# Simultaneously induced slow and fast gamma waves travel independently in primate primary visual cortex

**DOI:** 10.1101/2024.11.06.622198

**Authors:** Bhargava Gautham, Supratim Ray

## Abstract

Travelling waves have been reported for multiple neuronal oscillations across the cortex. Within the primary visual cortex (V1), large visual stimuli (gratings) can induce two gamma oscillations – slow (20-35 Hz) and fast (40-60 Hz), potentially due to two different interneuronal classes. However, wave like behaviour of these rhythms and their relationship is unknown. We showed large gratings to monkeys that simultaneously induced slow and fast gamma while recording from V1 using microelectrode arrays. Both slow and fast gamma were organized as travelling waves that were not locked to stimulus onset. Direction of wave propagation was significantly different but showed no correlation, even in concurrent waves. Slow gamma waves lasted for longer durations than fast gamma. Wave direction varied with stimulus orientation and spatial frequency. Thus, slow and fast gamma oscillations in V1 show unique spatial and temporal wave dynamics signifying that these gamma rhythms are generated by distinct neural circuits.

## Introduction

Neuronal oscillations are pervasive features of the mammalian cortex that have long been considered as candidates that aid in information transfer between different neural populations^1^. Dynamic synchronization of different neural populations could aid in diverse cognitive responses^2,3^. One method for coordination of oscillations are through generation of travelling waves (TWs), which are waves of propagating activity across patches of brain tissue that show a progressive temporal offset as distance from the point of origin increases^4^. Such TWs, initially observed in cortical activity of anaesthetized animals^5^, have now been established in awake states in primates^6^ and humans^7^, in the motor area^8^, visual regions such as the primary visual cortex (V1^6^) and area V4^9^, and in pathological conditions such as epilepsy^10^. While a clear physiological role is still being investigated for TWs, by virtue of their property to rapidly reorganize large areas of the cortex, they have been implicated in synchronizing neuronal populations, driving functional connectivity between neurons^11^ and shaping synaptic plasticity^12^.

Both TWs^6,9,13,14^ and neuronal oscillations^15,16^ have been extensively studied in V1 due to its ease of access, and vertical and laminar organization^17^ that serve as a template to model both these phenomena. While short and long range association fibres have been suggested to orchestrate similar synchrony in macroscopic recordings^18^, horizontal fibres in the lamina extending through layers 2/3 of V1 region of the occipital cortex have been implicated in the generation of TWs whose propagation speeds match the axonal conduction speeds in these unmyelinated fibres^19^ in microelectrode arrays. Local connectivity in V1 consisting of inhibitory and excitatory neurons, on the other hand, act in tandem in a delicate balance to give rise to neuronal oscillatory activity, such as the stimulus-induced narrowband gamma oscillations (30-70Hz), which have been implicated in a multitude of physiological functions including visual attention^20^, working memory^21^ and ageing^22^.

These stimulus-induced gamma rhythms can be consistently induced in the visual cortex by presenting gratings or bars. On presentation of sufficiently large gratings occupying more than 4° of the visual field, two distinct oscillatory activity have been observed in the frequency response of V1, termed as slow gamma (SG) and fast gamma (FG) in the range of 20-35Hz and 40-60Hz, respectively^16^, which have distinct tuning preferences over size, contrast, orientation and spatial frequency of visual stimulus showing that they represent different spatial extents of information coding. These two gamma rhythms are thought to be shaped by distinct inhibitory GABAergic subpopulations of interneurons^23–25^ with SG mediated through dendritic inhibition of the somatostatin interneurons and FG mediated via somatic inhibition of fast acting parvalbumin inhibitory neurons, albeit experimental evidence is mainly available from rodent studies.

Interestingly, several recent studies have suggested that the different interneuronal classes interact to modulate neural processing^26,27^. Therefore, if these interneuronal classes generate TWs, potential relationships between these waves (for example, direction of travel) could convey clues about their interactions. However, gamma band synchronization via TWs have been examined mainly during spontaneous activity, evoked responses or natural movie presentation in pre-specified bands^6,13,14,28^. Further, TWs are more prominent for small stimuli^29,30^, so it is unknown whether large gratings also produce SG and FG TWs, and whether these TWs potentially interact. We therefore recorded from V1 of monkeys using microelectrode arrays while presenting large gratings optimized to produce both SG and FG and studied potential wave propagation and interactions.

## Results

We recorded LFP from V1 area of two female monkeys while they viewed full screen sinusoidal gratings of varying orientations and spatial frequencies^16,31^. All analyses were done on 65 and 17 electrodes from the two monkeys for which stable receptive fields were obtained (see Methods for details).

### Characteristics of Slow and fast gamma oscillations

Large stimuli produced two gamma oscillations, whose power varied with orientation and spatial frequency (see Refs 16 and 31 for a detailed characterization). For Monkey 1, SG was strongest at 0° orientation while FG was strongest at 90° orientation. Figure 1A shows the change in power spectrum for trials when an orientation of 67.5° was presented (with spatial frequency of 1 cycle per degree (CPD)), which produced both gamma waves, as shown previously^16^. Gamma oscillations are known to occur in short bursts^32^, which is not evident in trial averaged spectra (Figure 1A) but are readily observed on individual trials (Figure 1B). To find gamma bursts on an individual trial, we filtered the signals in the SG (Figure 1C) and FG (Figure 1D) bands and identified the bursts using Hilbert Transform and thresholding^21,32^ (see Methods for details), as indicated by the black lines that correspond to high-energy patches in the time-frequency spectrum (Figure 1B). Figure 1E and 1F show these burst epochs for all electrodes in this monkey for that trial (the electrode that was shown in 1C-D is highlighted in dark blue). We found that gamma bursts tended to co-occur across multiple electrodes, as evidenced by aligned line segments in Figure 1E and 1F. We computed the fraction of electrodes that contained bursts within a trial (Figure 1G and 1H) and designated epochs where this fraction exceeded a threshold of 0.5 (dotted black lines) as gamma bursts for the entire electrode array (highlighted by thick horizontal lines in the bottom panels of 1G and 1H).

**Figure 1.**
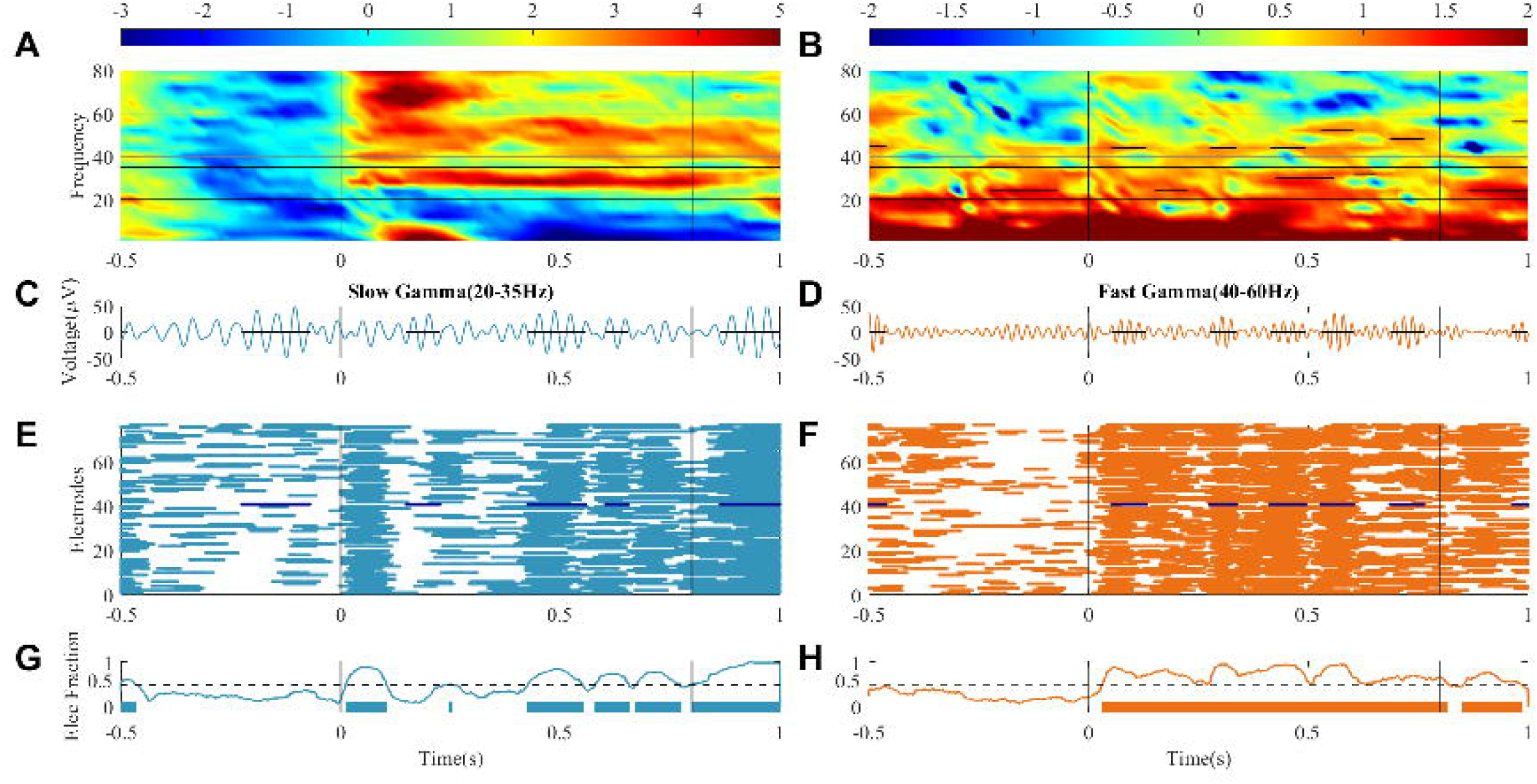
Analysis pipeline of the study. (A) Change in time frequency spectra averaged across all trials for one electrode in M1 when a full screen, 100% contrast stimulus of orientation of 67.5° and spatial frequency of 1cpd was shown for 800 ms (stimulus period indicated by vertical black lines). This stimulus combination was chosen because it induced strong gamma oscillations in SG (20-35Hz) and FG (40-60Hz) bands, represented by the thin horizontal black lines. (B) Raw time-frequency spectrum for a single trial for the same stimulus and electrode as in A. The interrupted horizontal black lines represent the gamma bursts calculated using Hilbert transform, as described below. (C-D) Gamma bursts were identified by filtering the signal in SG (C) and FG (D) bands and identifying epochs of high power. The plots show the filtered signals in blue (SG) and orange (FG) and identified bursts in black. These black line segments are also shown in B. (E-F) Gamma bursts for all electrodes for the same trial that was used in B-D. The electrode used in B-D is highlighted in dark blue. (G-H) Electrode fraction time series which indicates the fraction of electrodes in the array that contained a burst at any given time. Time points at which this fraction exceeded 50% (dotted horizontal line) were considered a burst for the entire array (as shown in the bottom of the plots) and were used for subsequent traveling wave analysis.

### Characteristics of Slow and fast gamma traveling waves

Within these epochs, TWs were defined as periods with sequential changes in phase calculated by circular linear correlation between the instantaneous phase of a given electrode with its position in the grid. This phase gradient was quantified by the phase gradient directionality (PGD), a derivative of the circular linear correlation coefficient and a direction measure that gives the direction of propagation of the wave (see Methods for details). We thus identified TWs for both SG and FG. A single TW was defined as a time period during which the direction of the wave did not change by more than 10° across successive time points (other criteria also gave similar results, as discussed below). We found that these large stimuli indeed produced TWs in both SG and FG bands (Figure 2 and Supplementary video 1). Of the TWs in SG and FG, overlapping TWs were defined as waves in the SG and FG bands during the same trial which had ≥50% of the same time points. Figure 2A shows the segmented waves in 20 trials where overlapping waves were observed along with waves in SG and FG that did not show any overlap in M1. TWs in SG and FG were thus observed to occur both independently and concurrently in different epochs.

**Figure 2.**
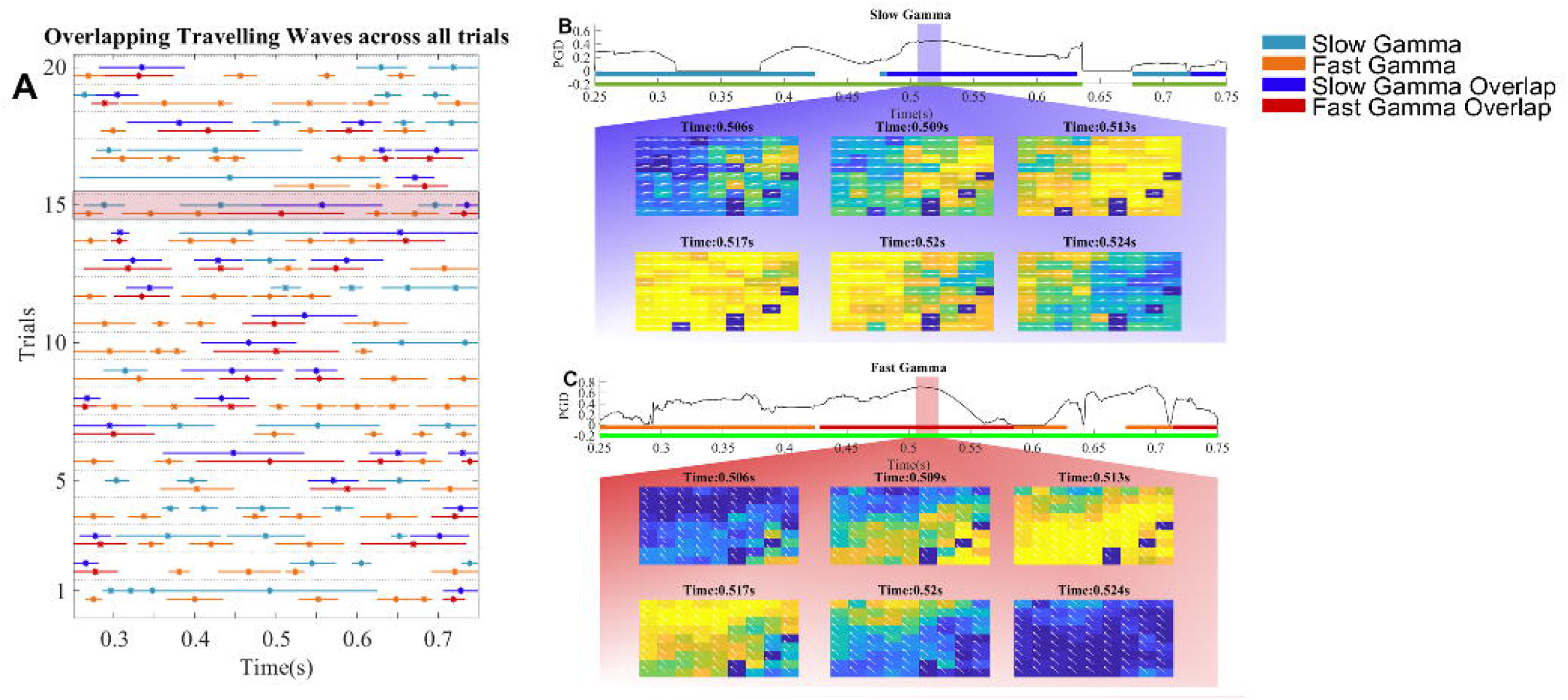
TWs were identified in the stimulus window of 0.25-0.75s across all trials for both SG and FG bands. This time period was used to avoid the initial transient after stimulus onset. Individual waves in SG and FG were considered to be overlapping if ≥50% of time points overlapped. (A) TWs of SG and FG for different trials, both overlapping (dark colors) and non-overlapping (light colors). B) A sample SG TW selected from trial 6, as highlighted by the red box in A. The selected wave has been highlighted in blue accompanied by the PGD values across the trial. Phase values across the grid were plotted using pseudocolor for the overlapping time points and direction of wave indicated by white arrows. C) A wave in FG occurring at the same time as the SG wave in B, with corresponding PGD values. B and C together show distinct phase propagation patterns and directions at the same time points for SG and FG.

Waves visualized in the trial highlighted in red in figure 2A have been expanded in Figure 2B and 2C and also shown in Supplementary Video 1. Figure 2B and 2C show simultaneously induced SG and FG waves whose directions were 208.16° (SG) and 271.47° (FG) during the overlapping period. Thus, overlapping waves in SG and FG, qualitatively, showed different propagation directions.

To quantify propagation directions in SG and FG, we plotted polar histograms of the direction distributions for M1 and M2 (figure 3). Directions of all waves (both overlapping and non-overlapping) for M1 in SG and FG bands are shown in Figure 3A while the mean angle is indicated by the colour coded arrows. Here, the direction is computed for a given wave by taking the circular mean across the period of the wave (as described in methods). TWs in the SG and FG bands propagated along mean directions of 230.73° and 239.42° and overlapping TWs (Figure 3B) propagated along the directions 230.91° and 250.12°. The mean directions for this monkey were not significantly different between SG and FG (p=0.589, Watsons U2=0.0709, Watsons U2 permutation test). For M2, mean directions were 77.97° and 117.65° for all waves (Figure 3D) and 48.38° and 118.96° for overlapping waves (Figure 3E), which were significantly different (p=0.002, Watsons U2=0.3078, Watsons U2 permutation test). All direction distributions were highly non-uniform (less than p<0.001 for all conditions; Hodges-Ajne’s^33^ test for non-uniformity).

**Figure 3.**
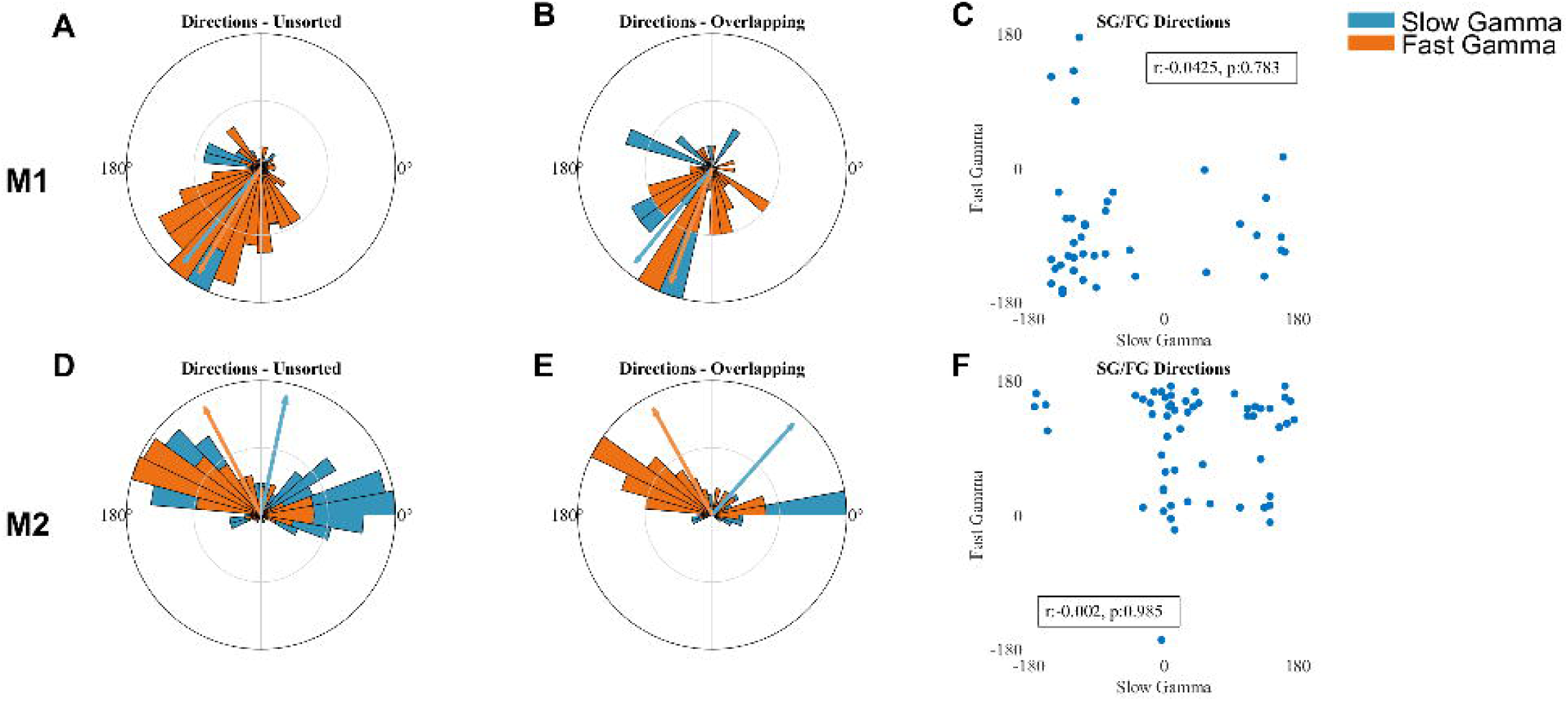
A) Direction distributions of all SG (blue) and FG (orange) TWs across all trials plotted on polar axis as a histogram for MI. Means of these directions are indicated by arrows. B) Same as A but for overlapping TWs. C) circular correlation coefficient between SG and FG directions for overlapping waves. Correlation coefficients and p-values are indicated in the plot. D-F) Same as A-C for M2.

These results show that SG and FG waves did not move randomly. The key question is that when generated simultaneously, whether the overlapping waves had any relationship. To address this, we plotted the directions of SG versus FG for the overlapping waves (Figure 3C and 3F) and employed circular correlation (CC) to test for correlations in these scatter plots. Importantly, the CC was not significant for either monkey (p-values indicated in the plots), suggesting no interaction between the two waves.

In the preceding analysis, we defined TWs as epochs where the wave direction did not change abruptly (maximum change of 10° between successive time points), but this criterion allowed a single wave to have slowly varying directions. In such conditions, taking the mean direction over the entire duration of the wave may not be informative. To address this, we also tested for wave similarity using a more stringent criteria where only a 5° variation throughout the wave was considered. In this procedure as well, circular correlation showed no significant relationship (M1: r = 0.208, p = 0.468 and M2: r= −0.009, p=0.968, by circular correlation^33^). As a final control, we also computed circular correlations between the directions at individual overlapping time points (instead of taking the mean directions), which also showed similar results (M1: r=-0.039, p= 0.011 and M2: r=0.003, p=0.813).

### TW properties in SG and FG bands

Apart from the propagation direction, we characterized TWs by their speed (in cm/s) and duration (in ms). Mean (± 1SD) propagation speed of the waves was 29.09±5.04 cm/s for SG and 27.71±13.86 cm/s for FG in M1 and 29.09±5.04 cm/s for SG and 27.71±13.86 cm/s for FG respectively in M2. No significant difference was observed between SG and FG (p>0.05 for all comparisons; Mann Whitney U test), however, FG showed larger variance in speed compared to SG.

A single wave was considered between two time points with a nonsignificant PGD value. The distribution of durations showed was skewed towards a non-normal distribution as seen with the durations of gamma bursts in each electrode. Mean (± 1SD) wave duration was 152.06±134.21ms (median=115ms) and 115.08±67.08 ms (median=101ms) in SG and FG in M1, and 90.87±41.23 ms (median=88ms) and 84.67±41.73ms (median=76 ms) in M2. Durations of SG were significantly higher than FG in M1 (p=0.03, Z=1.839, Mann Whitney U test) and in M2 (p=0.03, Z=1.812, Mann Whitney U test).

### Comparison of TW across multiple orientations and spatial frequencies

The analysis so far was restricted to a single orientation (67.5°) and spatial frequency (1 CPD) for which both SG and FG were strongly induced. We next tested whether the wave directions depended on stimulus properties. Figure 4A and 4B shows the circular mean of the direction distributions for each orientation (at a spatial frequency of 1 CPD) for SG and FG for M1, while Figure 4D and 4E shows the same for M2. Visually, we observed differences in direction distributions in both SG and FG across orientations. A multi sample circular analogue of Kruskal Wallis (Watson-Williams test^33^) was used to compare the directions, which revealed significant differences in directions in SG (p<0.001, F=4.42) and FG (p<0.001, F=10.83) for M1 and in SG (p=0.004, F=2.93) and FG (p<0.001, F=6.28) in M2.

**Figure 4.**
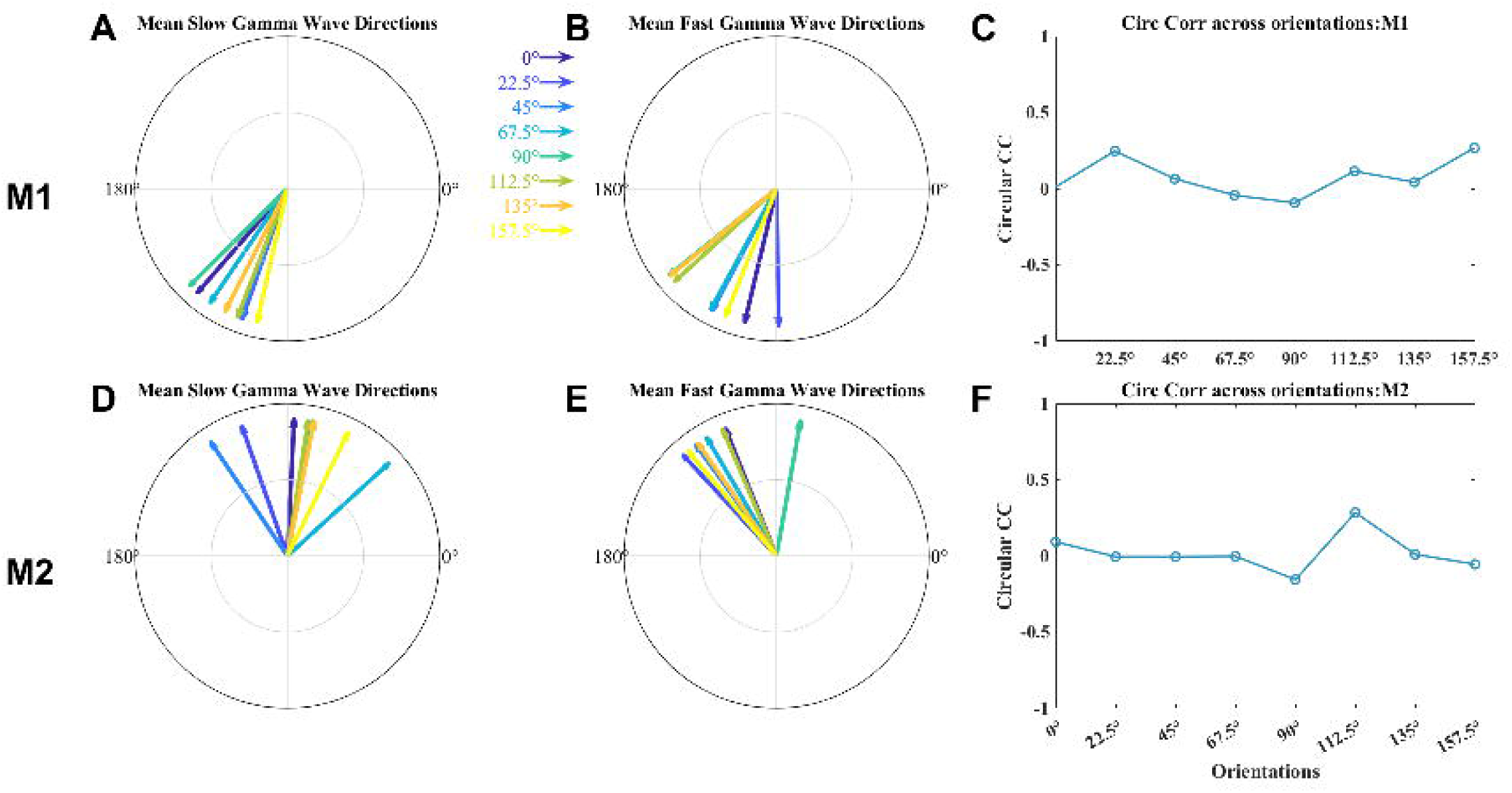
A-B) Mean directions of overlapping TWs for different orientations for SG (A) and FG (B) for M1. C) Circular correlation between SG and FG for different orientations, which were not significant for any orientation. D-F) Same as A-C for M2.

However, no consistent pattern in the change of direction was observed across both subjects. We also tested for correlations between overlapping SG and FG waves using circular correlation for each orientation (Fig 4C and 4F), but these correlations were small and not significant for any orientation (p>0.05 for all comparisons).

Similar results were observed when we tested for different spatial frequencies while keeping the orientation fixed at 67.5°: the mean wave directions varied significantly with spatial frequency but did not show any systematic pattern, while the circular correlation between overlapping SG and FG directions were small and not significant for any spatial frequency (data not shown).

### TWs in relation to gamma bursts

Finally, we investigated the temporal relationship between gamma bursts and occurrence of TWs within those bursts. Gamma burst in this context refers to the time segments during which more than half the grid had gamma activity, referred to as electrode fraction (as detailed in methods). To test when TWs appeared within the bursts, the burst segments were normalized to unit length. Figure 5A-B shows the location of the TWs within each of these normalized bursts for SG (Figure 5A) and FG (Figure 5B) for M1. We then binned this duration into 20 segments and computed the fraction of bursts that contained a TW at each of the bins and normalized the counts across the bins (termed as wave fraction, Figure 5C). Visually, it was observed that TWs were not consistently active throughout the duration of the burst fraction for both gamma bands. Incidence of TWs was greater in the centre of the gamma bursts compared to the initiation or termination of the bursts suggesting that TWs build up and drop off slowly once the gamma burst is generated. Similar results were obtained for M2 (Figure 5E-G). To quantify this result, a Kruskal Wallis test across TW probabilities for each bin was performed. Bins 1 and 20 had significantly different mean ranks compared to bins 2-19, computed for all trials for both subjects (M1: SG, p<0.001, F=4.42 and FG, p<0.001, F=10.83. M2: SG, p=0.004, F=2.93 and FG, p<0.001, F=6.28) across different orientations and different spatial frequencies (M1: SG, p<0.001, F=7.80 and FG, p<0.001, F=12.58. M2: SG, p=0.002, F=5.63 and FG, p<0.001, F=7.13). We extended this for comparison across all orientations and observed similar results (Fig 5D and 5H).

**Figure 5.**
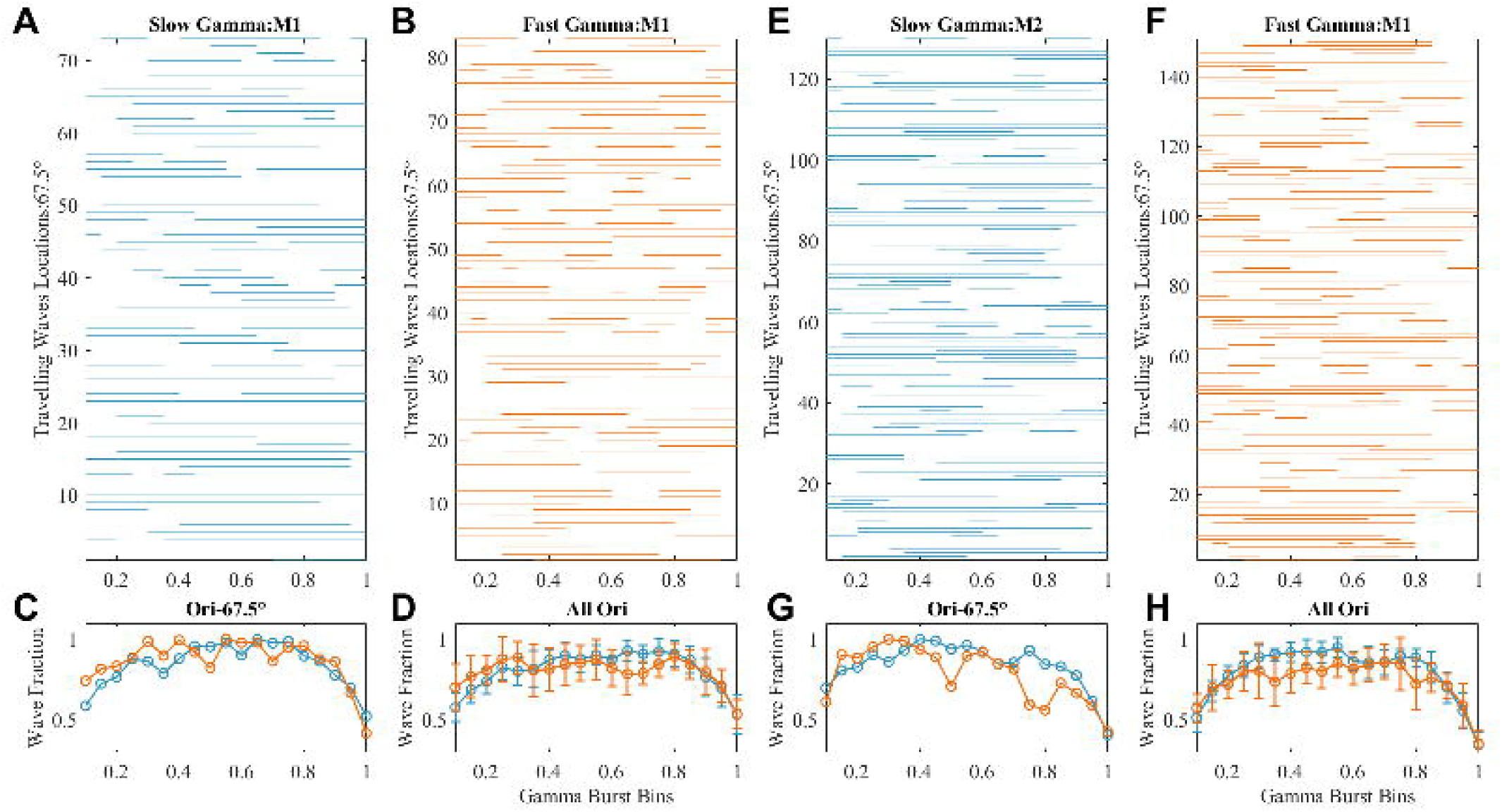
Temporal evolution of waves within bursts. A) SG bursts in M1 with a duration ≥50ms for the stimulus condition shown in Figure 1-3, normalized to 0-1 seconds and binned with a spacing of 0.05. The lines indicate the presence of a TW within this burst. B) Same as A but for FG. C) Fraction of bursts containing a TW at each of the time bins, termed as wave fraction in orientation of 67.5 °. D) Same as C, but after averaged across all orientations. Error bars indicate standard error of the mean. E-H) Same as A-D but for M2.

## Discussion

We show that both SG (20-35Hz) and FG (40-60Hz) oscillations in V1 induced by presentation of large static gratings were organized as TWs, which moved at non-myelinated nerve conduction velocities of ∼0.3m/s. TWs for SG and FG waves displayed unique propagation patterns evidenced by the direction distribution of the waves. Of the fraction of waves that showed overlap during the same trial, directions of SG and FG waves were non-correlated, indicating that distinct networks are employed for transmission of waves for both gamma rhythms. TWs were longer for SG than FG. This behaviour was preserved across multiple orientation and spatial frequency combinations.

Along with burst detection, we used an additional measure, electrode fraction, which ensured that wave detection was carried out only periods of high gamma power across at least 50% of the grid. However, mean wave size indicates that TWs consistently occupied >65% of the grid in both M1 and M2 (data not shown). Importantly, our method gives a dynamic measure to chart change in gamma power across the grid compared to earlier methods where electrode clusters tested for waves were fixed for each trial^7,8^.

### Mechanisms underlying generation of gamma oscillations and TWs

TWs have been postulated as a mechanism of neural synchronization across large patches of cortex^9^. In the visual cortex, TWs have been identified in the alpha^34^ and gamma bands^6,28^. Such propagating waves require tight knit interactions of two sets of cortical circuitries, one for the generation of oscillatory activity and another circuit to determine the spatial spread of the wave.

#### Generation of oscillatory activity

Of interest in this study was wave behaviour in different bands of gamma rhythms, namely SG and FG. The presence of multiple gamma rhythms has been identified in various cognitive domains such as memory consolidation in areas such as the hippocampus^35^, olfactory bulb^36^, entorhinal^37^ and visual cortex^16^. Within V1, the two gamma rhythms show different tuning characteristics^16^ suggestive of separate functional significance and networks. SG and FG in our study could be orchestrated through different inhibitory interneuron populations, namely somatostatin (SOM) and Parvalbumin (PV) interneurons. These interneurons vary with respect to the size of their synapses, locations, and their connectivity patterns^38^. PV interneurons are incident on the soma and are involved in supressing fast sodium currents and SOM interneurons show dendritic inhibition and supress slow dendritic calcium spikes, which can lead to different spiking patterns^38^.

#### Generation of TW

An important aspect in determining the spatial spread of the wave is the distance dependant delays between from the onset of the oscillation determining the amplitude decay of the wave, which is governed by the horizontal fibre connections^39^. Long range horizontal connections from excitatory pyramidal neurons that extend over several millimetres^40^ and have been postulated to form preferential connections with iso-feature selective columns^41^. While horizontal fibres do not confer specificity to the propagation direction of a wave and other wave properties as they lack any modulatory input, GABA-ergic inputs to pyramidal cells have been considered to curtail the radius of the wave and impart context sensitivity to the wave^42^. Computational studies have shown that generation of TWs require a non-homogenous population of cells, i.e., an interacting excitatory and inhibitory network^43^. In our data, direction distributions of SG and FG showed a significant difference, indicating that independent waves of both rhythms follow unique propagation pathways in both subjects. Additionally, consistent differences in wave durations between SG and FG in both subjects also supports the dichotomy of two TW generation circuits. Distinct connectivity profiles of PV and SOM interneurons may be responsible for the differential properties of TWs during propagation.

#### Potential role of TWs

An ongoing wave modulates the spike timing of the surrounding neurons based on the temporal window of its oscillation^44^. It follows that wave characteristics will vary based on the content of the inciting oscillation. Such an organization of pyramidal cells, interneurons and horizontal fibres has given credence to TWs acting to bind distant components within the cortex. Within the visual cortex, stimulus sensitivity is organized vertically in cortical columns, where each column has a specific receptive field^45^ and communication between adjacent receptive fields is through horizontal connections by the excitatory pyramidal cells^46^.

### Propagation direction of TWs in SG and FG

Patterns in wave propagation were identified by the direction of the phase gradient across the grid at each time point within the gamma burst. Statistically significant phase gradients were identified by the wave strength (PGD). Wave direction is an important property of TWs that determines cortical path along which the wave recruits neural populations. Wave direction represents the sequential reorganization of neural substrates along the gradient of its propagation^47^. Therefore, unique directions give rise to activation of different upstream ensembles and distinct perceptual or behavioural responses, such as posterior to anterior directions during memory encoding and vice versa during recall in the frontal lobe during an episodic memory task^47^. While it is not yet known if TWs are purely an epiphenomenon of the underlying oscillation and its circuitry or serve a unique functional purpose^6^, nevertheless, wave direction has been implicated in cognitive processes^47^.

TWs observed in our study were not locked to stimulus onset, neither were the gamma bursts, which has been observed previously^6^. The wave directions, however, were consistent across trials for a given stimulus combination (orientation and spatial frequency). While dynamics of gamma is preserved across species, underlying functional connectivity is highly individualistic. It might be prudent to comment on the substrates on which SG and FG TW is incident on in recordings from larger arrays or whole head recordings, which is not possible from this present study.

### Effect of stimulus parameters on TW direction

Gamma band in V1 has specific stimulus selectivity to features such as colour, size, contrast, orientation, and spatial frequency. In our study, the stimulus size and contrast were constant while orientation and spatial frequency was varied across trials. Histochemical studies have shown a patchy representation of neurons that are tuned to specific orientations separated by blobs in the striate cortex^48^. Hubel and Weisel^17^ have described a vertical organization of excitatory and inhibitory cells in V1 that respond to distinct stimulus features and synchrony was considered between columns of similar stimulus sensitivity for ensemble processing, although TWs that span across multiple such columns may seem counter intuitive to this kind of functional organization leading to averaging out of the stimulus specific information. However, in such an organization, the pattern of incoming excitation will change depending on stimulus properties, which could explain the differences in direction distributions for different orientations and spatial frequencies as observed in our data.

TW occurrence has been inversely associated with stimulus contrast and size^29,30^. The presence of both, high contrast and large stimulus size seem to reduce wave activity. We had to use full screen gratings because robust SG requires activation of a large patch of cortex^16^. The mechanism of TW generation when such a large patch of cortex is activated is likely to be more complex than the case when a small patch is activated (e.g. for a small stimulus) and TW propagates through the horizontal connections. Future studies where more complex activation patterns such as rotational waves are considered for stimuli of varying sizes may shed light on TWs generated by large versus small stimuli.

### Relation of TWs and gamma bursts

In V1, gamma rhythms are sporadic and bursty^32^, implying a waxing and waning of gamma power over the duration of the burst instead of a constant throughput from oscillating networks. In our data, we observed that a statistically smaller number of TWs were distributed in the ends of the bins compared to the rest in both SG and FG (Fig 5C-D and 5G-H). We also observed that gamma power tended to be more in the middle of the burst than at the ends (data not shown). Therefore, TWs were more likely to occur as the gamma power across the grid increased. Such a relationship between oscillatory power and wave strength has been established in the insula, where wave strength varied with theta and beta oscillations^7^. As gamma rhythms modulate spiking activity across the neural population as per changes in their oscillation cycle^49^, similar to waves of motor cortex during spontaneous activity^44^, gamma bursts can thus potentially entrain surrounding neurons into an ongoing wave given that a sufficiently large population of neurons are engaged in the oscillation^43^, which is then maintained by weak modulation of the wavefront determining the distance travelled by the wave. It is likely that such a co-ordinated rise in power required to initiate a wave leads to trial-by-trial variation of wave occurrence, wherein waves are not tied to the stimulus onset. Whether these TWs play a role or are an epiphenomenon, their occurrence, direction, duration and interactions provide important clues about the underlying cortical circuitry.

## Methods

### Animal recordings

Two adult female bonnet monkeys (Macaca radiata, weights: 3.3kg and 4 kg) were used in this study. All experimental protocols were in accordance with the guidelines approved by the Institutional Animal Ethics Committee (IAEC) of the Indian Institute of Science (IISc) and the Committee for the Purpose of Control and Supervision of Experiments on Animals (CPCSEA). For both monkeys, a titanium head post was surgically implanted followed by a period of training for a visual fixation task. Once the animals were trained, we installed a hybrid array of microelectrodes (Utah array; Blackrock Microsystems) consisting of 81 microelectrodes (9×9) and 9 ECoG (3×3) electrodes in the primary visual cortex (V1) of the right cerebral hemisphere, under anaesthesia. Grid placement was roughly set at ∼15 mm rostral from the occipital ridge and ∼15 mm lateral from the midline in V1. The microelectrodes were 1 mm in length with an interelectrode distance of 400 μm. The reference wires were placed over the dura near the recording sites or wrapped around titanium screws on the surface of the skull near the craniotomy. The receptive fields of the neurons recorded from the microelectrodes were centred in the lower left quadrant of the visual field at an eccentricity of ∼3° to ∼4.5° in Monkey 1 (M1) and ∼1.4° to ∼1.75° in Monkey 2 (M2). As in our previous studies, only electrodes for which reliable estimates of receptive field centers were obtained and for which the impedances were between 250 and 2500 KΩ were selected for further analysis, yielding 65 electrodes for M1 and 17 electrodes for M2.

### Experimental setting and behavioural task

For the task, the monkeys were seated inside a Faraday cage enclosure (to shield from external electrical interference) in a chair with their head fixed by the head post. The visual fixation task performed by the macaques required them to hold their gaze within 2° of a small fixation spot shown at the center of a gamma-corrected LCD monitor screen. The monitor was placed 50cm away from their eyes ensuring that the stimuli (full screen gratings) covered approximately a width and height of 56° and 33° of visual field respectively.

At the start of each trial, the monkey was required to hold its gaze on a fixation spot. Following an initial blank period of 1000 ms, a series of two to three stimuli were displayed in succession for 800 ms each, with an interstimulus interval of 700 ms. On maintaining fixation throughout the trial, a reward of a drop of juice was given to the monkeys. The stimuli, which were full screen sinusoidal luminance gratings, were presented at full contrast at 1 of 5 spatial frequencies (0.5, 1, 2, 4, and 8 cycles per degree) and 8 orientations (0°, 22.5°, 45°, 67.5°, 90°, 112.5°, 135° and 157.5°). For M1, the average number of trials each orientation and spatial frequency condition was 33 (28-36) and for M2, it was 42 (37-45).

### Gamma burst detection

All analysis was carried out in MATLAB 2023b. Gamma bursts were identified by using instantaneous power extracted using Hilbert transform^21,32^. Gamma bursts were extracted for all trials in two frequency bands, SG (20-35Hz) and FG (40-60Hz). Signals were filtered between these ranges using a 4^th^ order butterworth filter in both directions to avoid phase distortion. The instantaneous amplitude extracted as the absolute value of the bandpass filtered analytical signal and squared to obtain the instantaneous power. A gamma burst was said to present during a single trial if the gamma power was greater than 3 times the median instantaneous power in baseline.

### Identification of TWs

TWs were identified by using the circular regression method which has been described in detail by Das et al^50^. Quantitatively, TWs were defined as a set of electrodes with consistent phase differences showing a stable gradient^7^. This systemic change in phase differences was identified by using circular linear regression wherein regression was carried out between the circular phase of electrodes involved and their respective position within the grid, given by the x_i_ and y_i_ coordinate. Instantaneous phase (θ_i_) was extracted by Hilbert transform after signal filtering (Butterworth filter, filter order – 4).

The circular linear fit was defined by

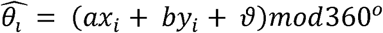

Where, θ^^^_i_ denotes the predicted phase, a and b are phase slopes corresponding to the rate of change of phase projected into each of the orthogonal dimensions and [ is the phase offset. Since the circular linear model does not have an analytical solution, an iterative fit was carried out. For this, the linear coordinates were first converted to polar coordinates, α = *atan2(b,a)*, and ξ = √*a^2^ + b^2^*, defined as angle of propagation and spatial frequency respectively. A grid search was conducted over and in step sizes of 1° and 0.5°/mm. Here, δ corresponds to the inter-electrode spacing, which was 0.4 mm. The model parameters and were fit to most closely match the observed phase at each time point. The goodness of fit (1), the mean vector length of the residuals between predicted and actual phases was defined as

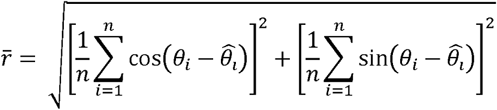

Where n is the number of electrodes. Values of α and ξ were chosen such that they maximize-1. To calculate statistical reliability, the phase variance explained by the best fitting model was calculated as the circular correlation ρ_cc_ between the predicted and actual phases.

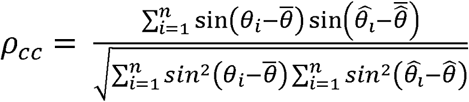

Where θ is the average of phases across the electrodes involved. To normalize ρ_cc_ as the number of electrodes involved can vary time point to time point,

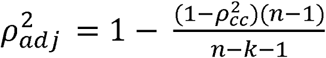

Where k is the number of independent regressors and *ρ^2^_adj_* is referred to as wave strength or phase gradient directionality (PGD). To characterize the features of TWs, we also looked at direction of propagation, wave speed (wavelength x frequency) where wavelength was taken as 2π/spatial frequency and duration of each traveling wave.

Thus, circular linear correlations in this analysis only considered the presence or absence of phase gradients, implying a travelling wave. We did not perform any classification of wave type, such as planar, radial etc. We also did not account for grid orientation to check if differences in direction distributions are accounted for between M1 and M2, therefore no anatomical correlates could be assigned to the direction distributions of the waves.

To test for statistical significance of wave strength, the linear coordinates were shuffled and PGD was re-estimated. This procedure was carried out 1000 times and only those wave strength values greater than the 95^th^ percentile of this distribution were taken as significant.

### Segmentation of TWs

TWs were considered to be present at time points where PGD values were significant beyond the permutation testing results. However, such segments may contain multiple successive waves, therefore, waves were selected that had a consistent direction of propagation across the grid with a variation of ±10° at each successive time point. Time points with variations greater than this threshold were part of a different wave. Thus, a time series of wave patterns was obtained for both SG and FG bursts.

We also used a more stringent criteria to segment waves, where a set of time points were considered to be part of a single wave if the mean variation of phase direction was only ±5° throughout the wave. Direction of propagation of each such wave was calculated by the circular mean of all direction values across all time points within a segmented wave. Such a procedure was adopted to enable comparison of wave direction between SG and FG rhythms using circular correlation. Circular correlation coefficient^33^ was used to calculate the similarity of directions between FG and SG waves. This was computed only between overlapping wave direction values as described above. We also used another approach where all overlapping time points between SG and FG, without considering the circular mean, were also taken up for circular correlation.

### Wave dynamics with respect to gamma bursts

To identify common behaviour across TWs across all trials, we collected gamma bursts across all electrodes were binned between 0 and 1 in steps of 0.05 to facilitate comparison. Only bursts greater than 50ms were used, similar to TWs. The probability of occurrence of a wave at a given bin was calculated as the fraction of bursts with a TW for that bin, termed wave fraction. The normalized mean of these histograms were plotted for qualitative comparison and a multi-sample circular analogue of Kruskal Wallis test^33^ was applied to test differences between distributions of wave probabilities across each bin for a template parameter combination (orientation: 67.5° and spatial frequency: 1CPD) and separately for trials of all parameter pairs.

---

**Supplementary Video 1:** Overlapping travelling waves from a single trial in M1 (as shown in Figure 2). PGD values in both SG and FG are shown and coloured bars below the abscissa represent wave times of SG and FG. Phase propagation patterns during intersecting time points in SG (left) and FG (right) show unique directions for each band.

